# Engineering bacterial protein polymers to support human pluripotent stem cell growth and differentiation in culture

**DOI:** 10.1101/2024.04.29.591606

**Authors:** Adam R. Creigh, Helen Waller, Jeremy H. Lakey, Zofia M. Chrzanowska-Lightowlers, Robert N. Lightowlers, Daniel T. Peters

## Abstract

Induced pluripotent stem cells (iPSCs) are of significant value due to their wide ranging potential, removing the need for embryonic material. To successfully culture, expand and differentiate these cells, it is crucial to maintain a precise biological environment, including an appropriate attachment substrate. Commonly used attachment substrates include recombinant extracellular matrix (ECM) components like vitronectin, as well as animal-derived ECM mixes such as GelTrex and Matrigel. However, there is growing interest in exploring alternative approaches to support bioactivity of cells. One approach that is gaining traction is the use of the Caf1 protein of *Yersinia pestis*. This protein is appealing primarily due to its stability, modularity, and ease of production. In this study, we have developed novel variants of Caf1 that effectively support the growth and differentiation of iPSCs, performing at least as well as GelTrex. Our findings highlight the potential of Caf1 laminin and vitronectin mimics as viable alternatives for supporting iPSC growth and differentiation. The successful development of these Caf1 variants opens new avenues for the field, paving the way for better defined, more cost-effective and readily available attachment substrates in iPSC research and applications.

## Introduction

The growth and differentiation of stem cells hold significant academic and commercial potential due to their ability to generate a wide range of cell types^1,2^. These cells can be utilized for various purposes, including research, as well as for clinical and non-clinical applications. These include creating patient-specific cells to investigate tissue specific diseases; with the use of mesenchymal stromal cells (MSCs) in autologous and allogeneic treatments for osteoarthritis^3^, and the development of chimeric antigen receptor T-cells (CAR-T) from induced pluripotent stem cells that could lead to an “off the shelf” allogeneic treatment for intractable cancers^4^. Additionally, there is growing economic interest in cultivating non-human stem cell cultures to generate muscle and fat cells necessary for cultivated meat production^5,6^.

Stem cells can be derived from various tissues at different developmental stages, including embryonic stem cells (ESCs) and adult stem cells (e.g., MSCs)^1^. Somatic cells can also be transformed into induced pluripotent stem cells (iPSCs) through the introduction of the key eponymous Yamanaka factors^2,7^. The use of iPSCs is advantageous compared to other cell lines due to their ability to differentiate into multiple cell types (pluripotency) without the need for embryonic material^8^.

To maintain their stemness and prevent the loss of pluripotency during differentiation into specific lineages, stem cells require a consistent biological environment that supports this state. Such conditions involve both the attachment substrate and a combination of defined growth factors added to the culture media. Typically, when growing these cells ex-vivo, extracellular matrix (ECM) proteins like laminin or vitronectin are used as components of the attachment substrate. These proteins are commercially available in the form of recombinant proteins. Alternatively, ECM mixtures obtained from animal sources, such as Matrigel or GelTrex, can be used. However, these can be costly, prone to batch-to-batch variation, and present challenges in practical use^9^.

In this study, our aim was to develop an alternative to these extracted ECM proteins, using an engineered polymeric bacterial protein, Caf1. Naturally produced by *Yersinia pestis*, Caf1 is composed of up to 250 non-covalently linked individual 15 kDa monomer subunits^10^ (**Fig. 1A**). The polymer is used to prevent recognition and phagocytosis of the pathogen by host macrophages, in effect forming a molecular “cloaking device” that surrounds the bacterial cell^11,12^. Previously we have shown Caf1 polymers can be produced recombinantly in *E. coli*, and used to coat cell culture surfaces, where in its natural form it retains its native properties, forming a Teflon-like “non-stick” surface that mammalian cells do not adhere to or interact with^13^. This allows bioactive peptide motifs, such as cell adhesion proteins of the ECM or growth factor molecules to be inserted into the Caf1 protein at various positions within the monomer structure (**Fig. 1B, C**). This provides us with the ability to exquisitely define what bioactive signals will be transmitted to cells (**Fig. 1D**). We have previously utilized this system to create functional mimics of ECM proteins such as fibronectin^13^ and laminin^14^, as well as growth factors such as BMP2^15^ and VEGF^14^.

**Figure 1:**
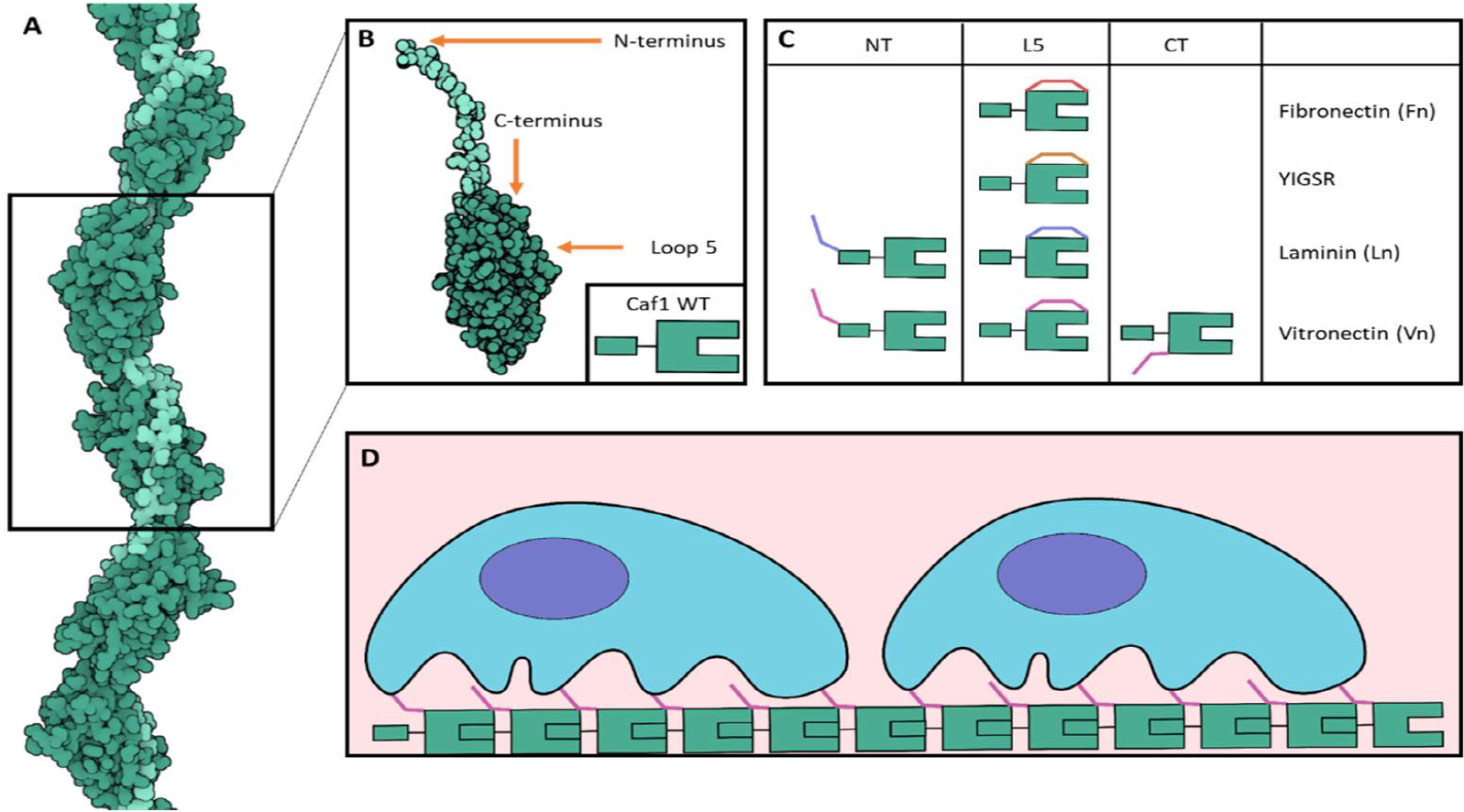
Cartoon depicting the Caf1 system used in this study. (**A**) Caf1 (green) forms modular, polymeric structures by donating its N-terminal donor strand (light green) to the next subunit in the chain to form a strong, non-covalent complex. (**B**) Overview of the Caf1 wild type (WT) subunit, with insertion points for bioactive motifs highlighted. (**C**) Overview of the modified Caf1 subunits used in this study. (**D**) Cartoon showing cells adhering to a Caf1-CT-Vn polymer.

Here, we test several variants of engineered Caf1 to find an effective substrate for iPSC attachment and proliferation. Caf1 was modified to incorporate key adhesive peptides of the structural matrix proteins fibronectin, vitronectin, and laminin. Two of the tested proteins, Caf1-CT-Vn, with a vitronectin sourced peptide, and Caf1-NT-Ln, containing a peptide from laminin, were able to successfully accomplish this task. The iPSCs grown on these Caf1 fusion protein matrices retained pluripotency markers and were able to freely form embryoid bodies that retained the capacity to differentiate into all three germ line lineages. We also show that human iPSCs grown on these chimeric Caf1 surfaces were able to successfully differentiate into beating cardiomyocytes when supplied with the relevant growth factors. Therefore, Caf1 multimers can successfully incorporate parts of both vitronectin and laminin to support the growth and differentiation of iPSCs. Combined with the other advantageous properties of Caf1, like the ability to mix and match multiple bioactive signals in a single material, this study provides an important step towards using Caf1 as a tool for culturing stem cells in both academic and industrial settings.

## Results

### Determination of Caf1 mimics that support iPSC attachment

A library of engineered bioactive Caf1 polymers has been produced over recent years that includes mimics of several ECM proteins^13–15^. To determine which of these variants could potentially support the attachment and growth of iPSCs, plates were coated with these Caf1 proteins. The variants (**Fig. 1C**) chosen included a fibronectin mimic (Caf1-L5-Fn), laminin mimics (Caf1-L5-YIGSR, Caf1-Ln variants) and vitronectin mimics (Caf1-Vn variants) where L5 indicates that the bioactive motif is inserted into loop 5 of the Caf1 protein. Caf1-L5-Fn and Caf1-L5-YIGSR have been shown to be pro-adhesive for other cell types in previous studies^13,14^. Caf1-Ln and Caf1-Vn had not been previously tested until this study and motifs were placed at both loop 5 and also the N-terminus (NT) of the protein. Caf1 wild-type (WT) was included as a negative control, as it is non-adherent for cells and the commercially available Geltrex (Gibco) was used as a positive control.

As expected, cells did not adhere to the Caf1-WT coating (**Fig. 2**), displaying a rounded phenotype. Caf1-L5-Fn and Caf1-L5-YIGSR also did not support cell adhesion. Of the Caf1-Ln and Caf1-Vn variants, only Caf1-NT-Ln showed a pro-adhesive phenotype, whilst cells adhered well to the Geltrex. A small number of elongated cells were present on Caf1-NT-Vn but did not go on to form a colony, unlike Caf1-NT-Ln and Geltrex. Therefore, a third insertion site for the bioactive vitronectin motif was tested, at the C-terminus (Caf1-CT-Vn). This protein was able to support cell adhesion and colony formation to a similar degree as both Caf1-L5-Ln and Geltrex.

**Figure 2.**
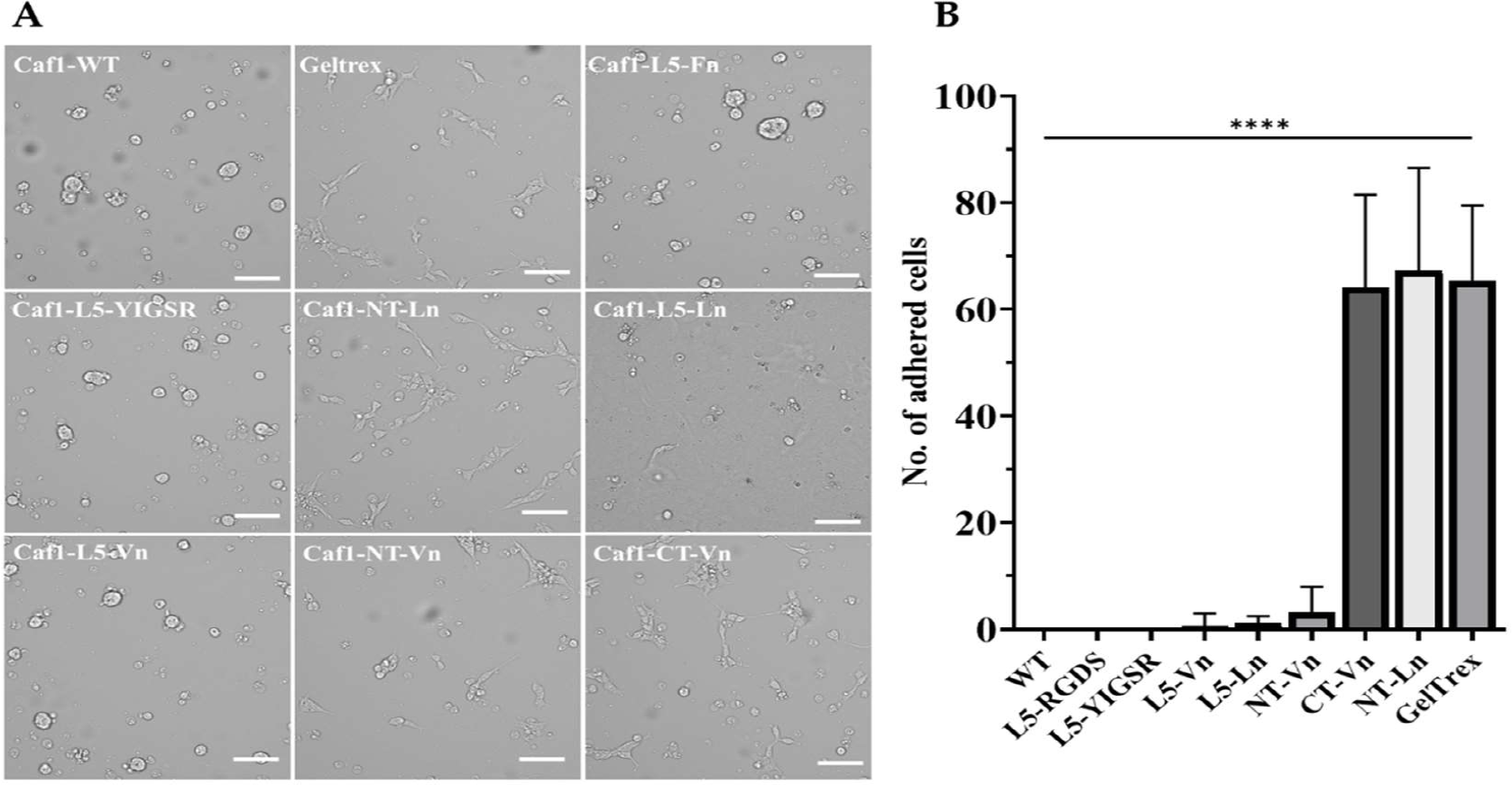
Evaluation of engineered Caf1 variants as suitable substrates for iPSC adherence. Plates were coated with several of the engineered bioactive Caf1 polymers containing ECM protein mimics in addition to Caf1 wild-type (WT) and Geltrex. iPSCs were seeded and incubated for 24 hours before imaging and cell counting per field of view. The images (**A**) represent twelve repeats per condition. Scale bar: 100 μm. Statistical analysis was performed on cell counts using one-way ANOVA, followed by Bonferroni’s multiple comparisons tests. **B** (n = 12 fields of view, \*\*\*\**p* < 0.0001). Abbreviations: L5, Loop 5; NT, N-terminus; CT, C-terminus; Fn, Fibronectin; Ln, Laminin; Vn, Vitronectin.

### Assessment of Caf1 adherent cultures for stemness markers

To ensure the Caf1 substrates were not negatively affecting the pluripotent nature of the iPSCs, we examined cultures after several passages on the substrates for retained pluripotency by immunofluorescence staining of key stem cell phenotypic markers (**Fig. 3**). Colonies grown on Geltrex, as well as Caf1-NT-Ln and Caf1-CT-Vn, were stained with antibodies recognizing OCT4, NANOG and TRA-1-60 and expression levels visualized via confocal fluorescence microscopy. In all cases, pluripotency markers were detected, and expression of Caf1 substrates appeared comparable to Geltrex, as expected for human pluripotent stem cells.

**Figure 3:**
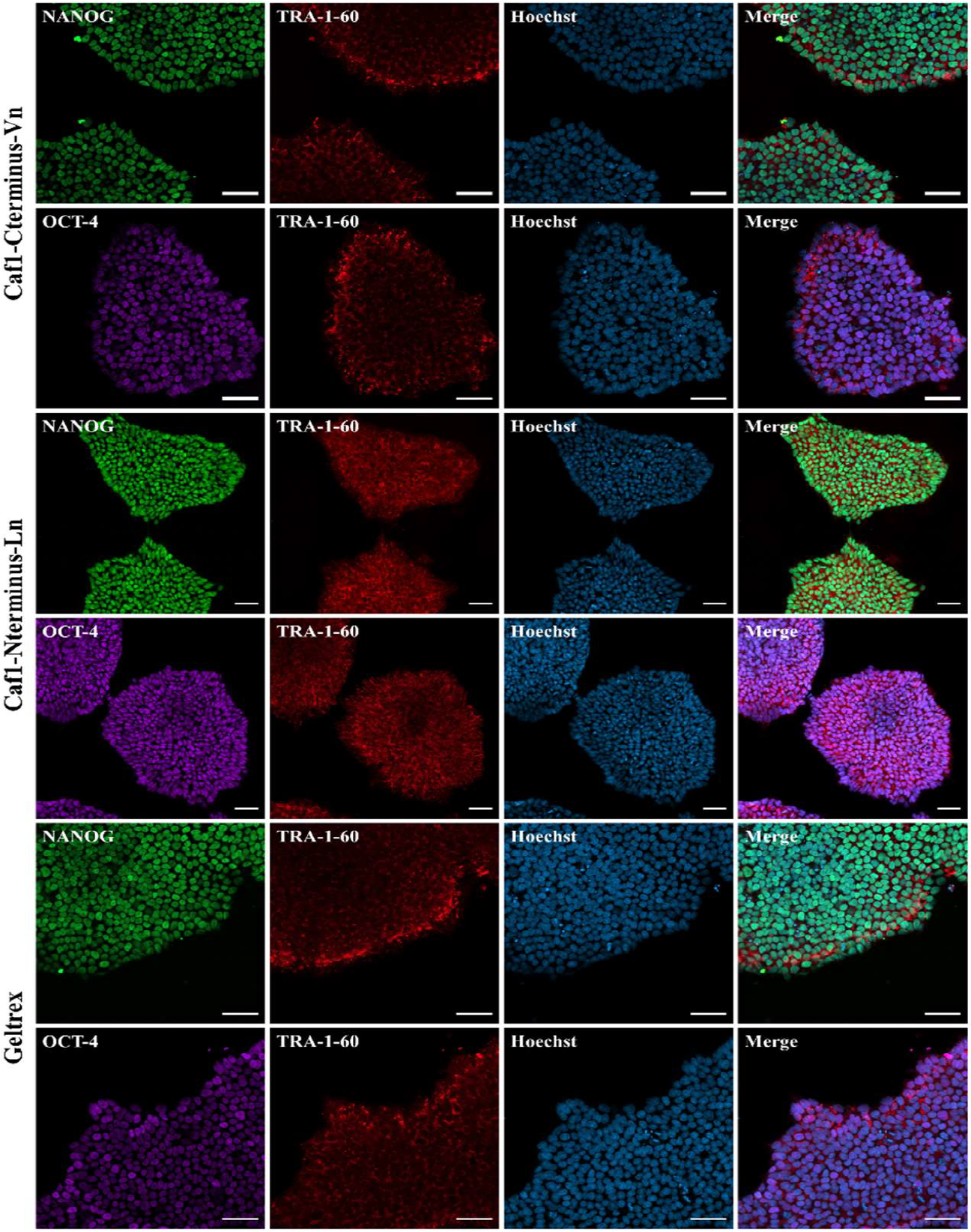
Immunocytochemical analysis of pluripotency markers of hiPSCs grown on Caf1 substrates compared to Geltrex. Cells were expanded over repeat subcultures before incubation with antibodies for transcription factors OCT4 (purple), NANOG (green), and surface antigen TRA1-60 (red) to highlight endogenous expression of pluripotency marker genes. Cultures were co-stained with Hoechst nuclear stain (blue) Scale bar: 50μm.

The gold standard test for evaluating pluripotency of iPSC cultures is to determine their ability to differentiate towards all three primary germ layers – endoderm, mesoderm, and ectoderm – which can be achieved by formation of embryoid bodies (EBs). iPSCs cultured on Caf1 substrates, as well as on Geltrex, over several sub-cultures were differentiated into embryoid bodies and visualised for expression of three key germ layer markers: nestin, α-fetoprotein, and smooth muscle actin (α-SMA) via Lattice-Lightsheet microscopy (**Fig. 4A**). EBs were a similar size and uniformly expressed all three markers. This was determined by gene expression analysis before and after differentiation via RT-qPCR using the TaqMan™ hPSC Scorecard™ Panel (**Fig. 4B**). In all cases, the cells grown on Caf1 performed similarly to those grown on Geltrex, and evidence of increased gene expression into all three lineages was observed. Therefore, Caf1-NT-Ln and Caf1-CT-Vn can act as effective substrates for iPSC adherence and proliferation, whilst maintaining the stem phenotype and preventing spontaneous differentiation, at least as effectively as Geltrex.

**Figure 4:**
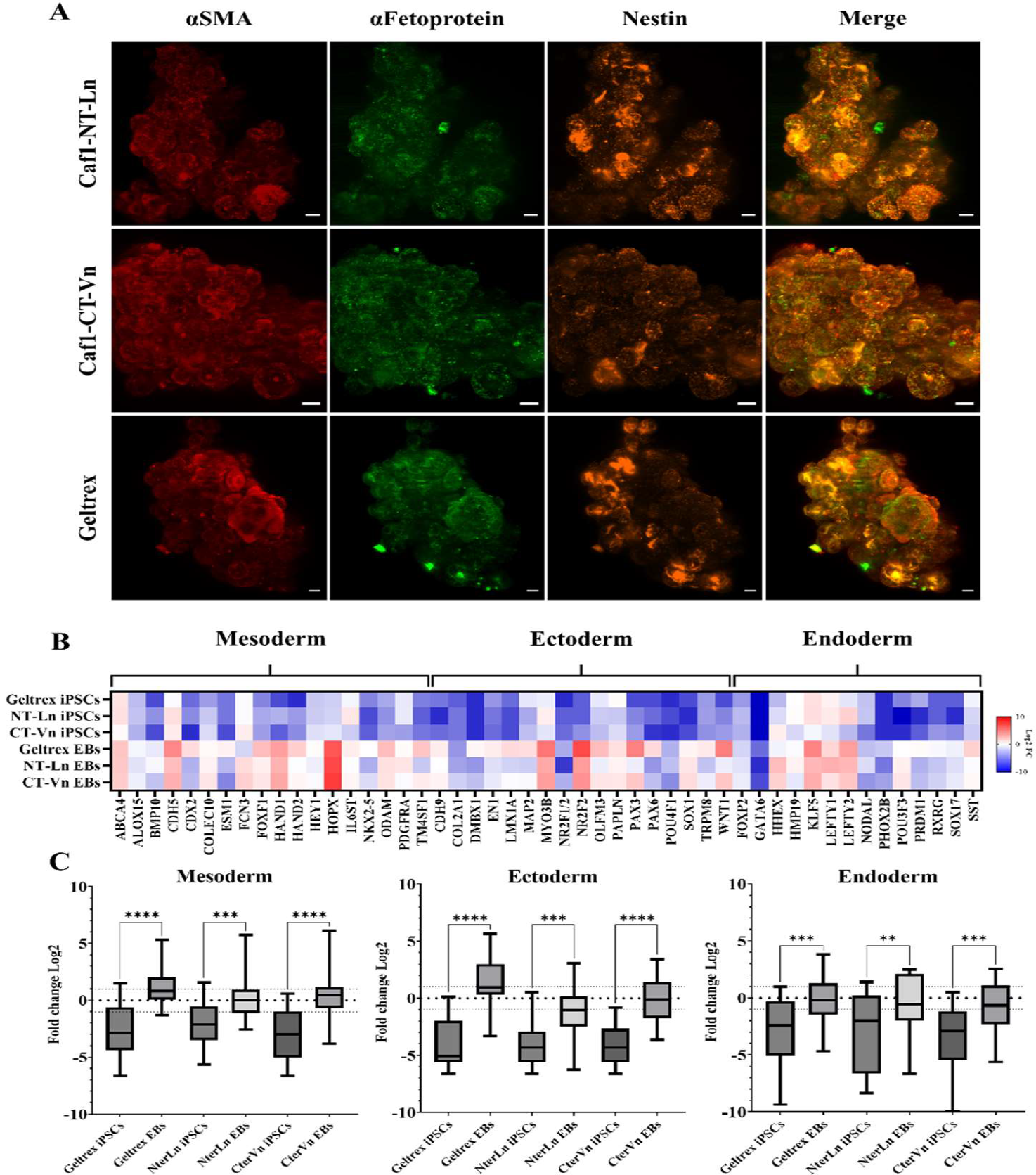
Embryoid body formation from iPSCs grown on Caf1 substrates and Geltrex. (**A**) Immunofluorescence analysis of 5-day embryoid body spheroids with markers of all three embryonic germ layers visualized using Lattice Lightsheet microscopy with antibodies for αSmooth muscle actin in red (Mesoderm), Nestin in orange (Ectoderm), and α-fetoprotein in green (Endoderm). *Cont*. Scale bars: 50μm. (**B**) Heat map of gene expression fold changes from TaqMan hiPSC scorecard assay with mesoderm, ectoderm, and endoderm gene markers highlighted. Color range illustrates reduced increasing expression (red), gene identifier shown below. (**C**) Boxplots correspond to center quartiles for all genes in each linage set, the mean is marked by a black bar and whiskers show total range of values. Significance between sets determined by one way ANOVA coupled with Šídák’s correction. Dotted lines plot baseline reference levels from supplier (<-1: downregulation, >+1: upregulation) plotted as Log_2_ of fold change. \*\*\*\**p* < 0.0001

### Caf1 supports the differentiation of iPSCs into cardiomyocytes

Next, to test whether Caf1 could support iPSCs during differentiation, iPSCs were grown on Caf1-NT-Ln or Geltrex before being differentiated towards cardiomyocytes using small molecule manipulation modified from Lian *et al* ^16^(**Fig. 5A**). On day 3, the morphology of the cells had changed and by day 14 clusters of beating cells could be observed. The cells grown on the Caf1 surfaces appeared morphologically similar to those grown on Geltrex (**Fig. 5B**). To determine successful maturation of cardiomyocytes, expression of the cardiac-specific marker cardiac muscle troponin T (cTNT) was probed by immunofluorescence (**Fig. 5C**). All contracting cells were positive for cTNT in the core of the cluster, where the beating cells were located. The cultures were also positive for α-SMA; an indicative marker of myofibroblast formation which also commonly arises during cardiac lineage differentiation. Therefore, Caf1-NT-Ln was shown to be able to support the attachment and differentiation of iPSCs.

**Figure 5:**
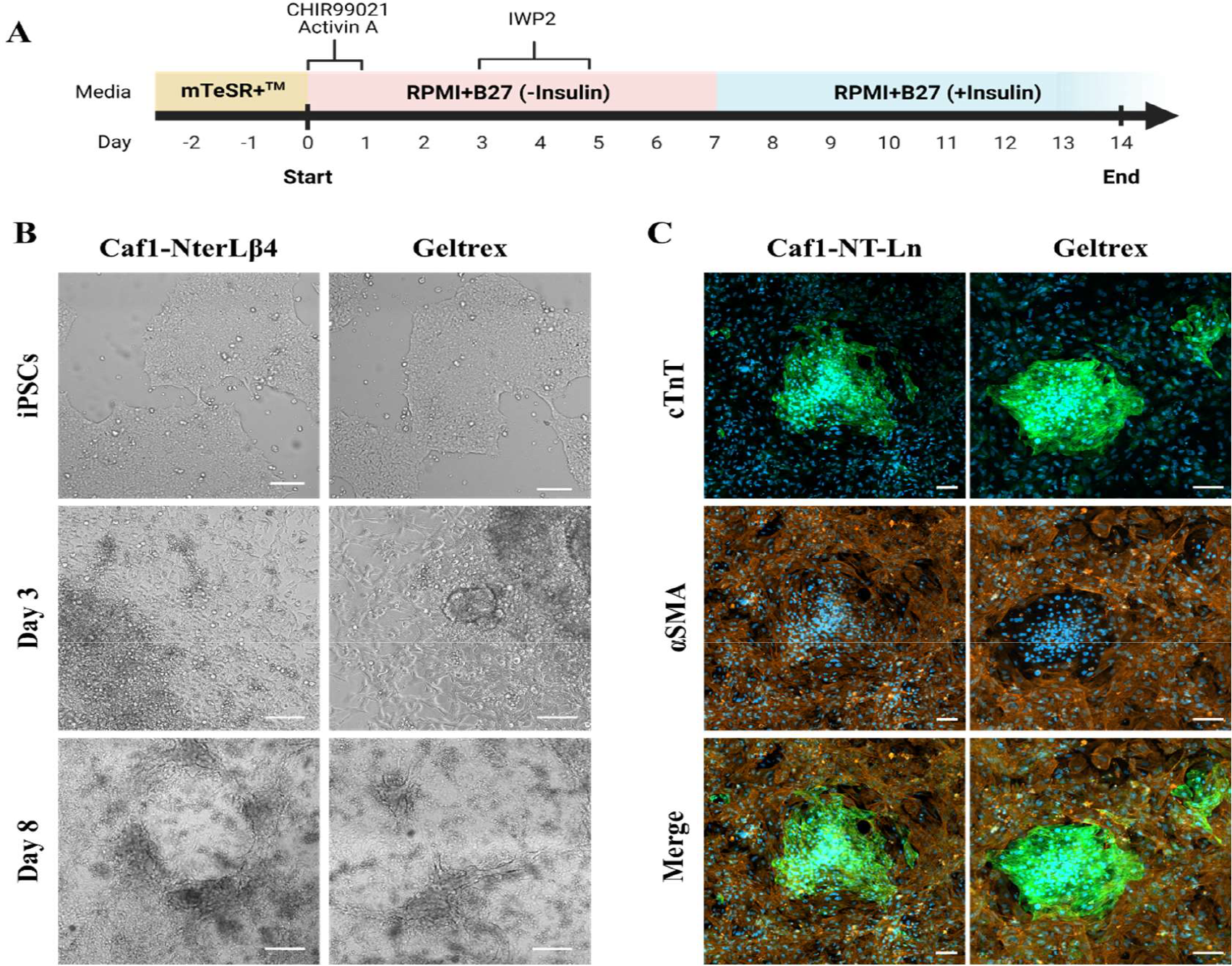
Differentiation of iPSCs into cardiomyocytes on Caf1-NT-Ln and Geltrex substrates. (**A**) Schematic of representation of small molecule differentiation protocol of iPSCs to contracting cardiomyocytes with media and small molecules defined at each stage. (**B**) Representation of the phenotypic changes in cell morphology seen throughout differentiation stages of both substrates. (**C**) Cardiomyocyte characterization via immunofluorescence analysis of cardiac linage specific surface markers; cTnT and α-SMA, stained on iPSCs differentiated to day 8 on Caf1-NT-Ln and Geltrex substrates. Cultures were co-stained with Hoechst nuclear stain. Scale bar represents 100μm.

## Discussion

The aim of this study was to develop variants of Caf1 that are able to mimic the ECM components (typically laminin and vitronectin^9^) that are commonly used to propagate cultures of iPSCs. The usefulness of Caf1 as a biomaterial is linked to a combination of several unusual properties that it possesses that have either been directly selected for, or are fortunate side-effects of the selected traits. Its native function is to protect *Yersinia pestis* bacteria from engulfment and subsequent destruction by macrophages upon entering a warm-blooded host^11,12^. It achieves this through three key methods: i) by acting as a polymer brush to lower the chance of macrophages interacting with it in the first instance, ii) by lacking any high-affinity adhesion sites for macrophage receptors, essentially making it “non-stick”, and iii) having very high mechanostability which is just sufficiently strong enough to resist the force that any macrophages that have adhered might exert onto the molecule, preventing it from breaking and revealing the bacterial surface (which contains a multitude of high-affinity targets for a macrophage)^12^. Hence, Caf1 is robust, modular, polymeric, and bioinert, but can also be expressed recombinantly in high yield by *E. coli* and can be modified to contain bioactive sequences^13,17^. This sets it apart from other proteins which, in general, tend to make poor materials on account of their low stability and low to moderate expression yields. Furthermore, we can also exploit Caf1’s non-covalent modularity through an *in vitro* refolding protocol that allows different subunits to be mixed together in a single, complex, multifunctional material^18^.

With these factors in mind, the systems in which Caf1 can be deployed to useful effect is increasing. For example, a recent study shows how Caf1 laminin and VEGF mimicking modules can be linked in a single polymer in order to facilitate the adhesion and migration of human endothelial cells within a 3D hydrogel network^14^. Another study shows the effect of Caf1 osteopontin/BMP2 mimicking mosaic polymers on primary human MSCs, triggering differentiation and the early stages of bone formation without the need for supplementary growth factors^15^. Thus, a range of Caf1 mimics have been developed including the above examples, as well as a fibronectin mimic^13^.

Interestingly, the previously developed fibronectin and laminin mimics tested here did not support the attachment and growth of iPSCs, leading to the development of two new Caf1 variants (Caf1-NT-Ln and Caf1-CT-Vn). In addition to the identity of the bioactive motif, the position of the motif within the Caf1 structure also played a major role in determining the bio-adhesive activity of the protein. For instance, the vitronectin motif worked well when placed at the C-terminus of Caf1, but was not effective at either the N-terminus or Loop 5 locus. This stands in contrast to other variants where these locations do have activity, e.g. the Caf1-NT-Ln shown here and also the Caf1-L5-Fn and Caf1-L5-YIGSR variants that have been shown to be bioactive with other cell types in previous studies^13,14^. These results are consistent with the view that, although a particular motif may be able to bind to a cell surface receptor, it must be presented in an accessible fashion and that this may differ for different motifs. It is therefore advantageous that the Caf1 subunit has several locations where motifs can be reliably inserted.

The results show that iPSCs grown on Caf1-NT-Ln or Caf1-CT-Vn surfaces retained both their pluripotency markers and their capacity to differentiate into different tissue types, including, as demonstrated, cardiac and smooth muscle cells. In this regard, the Caf1 surfaces could be positively compared to the commercially available product Geltrex, which is a basement membrane extract from animals that contains a mixture of ECM components, including laminin and collagen. Notably, as Caf1 is produced by bacteria it is free from animal components, and can also be used in a 2D monolayer as opposed to a 3D hydrogel, making it easier to use.

To conclude, this study demonstrates an important addition to the iPSC culture “toolkit” derived from Caf1 modules. We show the development of effective mimics of laminin and vitronectin that can be used to both grow iPSCs, and support them during differentiation protocols. In future work, these new modules could be combined in mosaic polymers with other versions of Caf1, such as those that mimic growth factors, to provide highly specific, multifunctional bioactive surfaces for the control of cell growth, differentiation and behavior.

## Materials and Methods

### Plasmids and Cloning

Sequences encoding Caf1-WT, Caf1-L5-Fn and Caf1-L5-YIGSR were present on pGEM-T plasmids prepared previously^13,14^. Sequences encoding Caf1-NT-Ln, Caf1-L5-Ln, Caf1-NT-Vn and Caf1-L5-Vn were produced by modifying pGEM-T Caf1-WT by PCR to include the bioactive peptide. The Caf1-CT-Vn gene sequence was synthesized and incorporated into a pET28a backbone by GENEWIZ™ by Azenta Life Sciences.

### Protein Expression and Purification

Caf1-WT, Caf1-L5-Fn, Caf1-L5-YIGSR, Caf1-NT-Ln, Caf1-L5-Ln, Caf1-NT-Vn and Caf1-L5-Vn were produced as described previously^13^. Briefly, *E. coli* cells transformed with the plasmid were grown at 35°C 180 rpm in shake flasks overnight using Terrific Broth (TB) media. Caf1 is exported from the cells by the Caf1M/Caf1A chaperone and accumulates in the media. The protein is then purified by tangential flow filtration using 500 kDa and 100 kDa MWCO filters, followed by size exclusion chromatography to remove any remaining impurities, with phosphate buffered saline (PBS) being the final buffer solution.

### Cell culture

Human dermal fibroblast-derived hiPSCs were cultured in feeder-free conditions on Geltrex™ LDEV-Free Reduced Growth Factor Basement Membrane Matrix (A1413201, Gibco) diluted 1:100 in ice cold Advanced DMEM/F-12 (12634010, Gibco) or modified Caf1 cell culture scaffolds coated 60 mm culture dishes with mTeSR™ Plus stabilised feeder-free maintenance medium (100-0276, STEMCELL Technologies) following the manufacturers’ recommendations. Cultures were dissociated by TrypLE (12605010, Gibco) and reseeded as a single cell suspension at a density of 1.5×10^4^ cells/cm in mTeSR™ Plus supplemented with 10 μM Y-27632 RHO/ROCK pathway inhibitor (72304, STEMCELL Technologies). Media was replaced with Y-27632 free mTeSR™ Plus after 24 hours.

### In vitro EB formation

Generation of embryoid bodies (EBs) was carried out by harvesting 80–85% confluent hiPSCs cultured in feeder-free conditions. Cells were dispersed into a single cell suspension, washed with DPBS and resuspended in complete EB medium, Knockout™ DMEM/F12 (12660-012, Gibco) supplemented with 20% KnockOut ™ Serum Replacement (10828-028, Gibco), 2 mM GlutaMAX™ (35050-061, Gibco), 0.1 mM MEM Non-Essential Amino acids (11140-050, Gibco) and 0.1 mM 2-Mercaptoethanol (31350-010, Gibco). Cell suspension/ EB medium is supplemented with 10 μM Y-27632 and seeded dropwise to a non-TC treated culture dish. Y-27632 is removed after 24 hours and media is refreshed every 48 hours.

### Immunofluorescence analyses

Immunostaining of hiPSCs was visualised via confocal microscopy (LSM800Airyscan, Zeiss) on cells that were seeded in glass-bottomed imaging plates (IBIDI). Cells were washed (DPBS, Gibco), and fixed with 4% PFA (for 10 min, at room temperature (RT)) then permeabilised and blocked in DPBS with 1% BSA, (w/v), 0.1% Triton X (v/v) for 1 hr at room temperature. All primary antibodies were diluted and incubated in DPBS/BSA/Triton solution overnight at 4°C. Cells were washed briefly in DPBS before the incubation with secondary antibodies and Hoechst 33342 nuclear stain for 1 hr at room temperature protected from light. Antibody suppliers and dilutions are as given: Primary NANOG (MA1-017, Invitrogen, 1:100), OCT-4 (701756, Invitrogen, 1:1000), TRA-1-60 Podocalyxin (14-8863-82, Invitrogen, 1:100); Secondary antibodies: Goat anti-Rabbit IgG Alexa Fluor 488 (A-11008, Invitrogen, 1:250), Goat anti-Mouse IgG1 Alexa Fluor 488 (A-21121 Invitrogen, 1:250), Goat anti-Mouse IgM (A-21238, Invitrogen, 1:250).

### Tri-lineage analysis using Quantitative RT-PCR

Total RNA was isolated from both undifferentiated hiPSCs and derived embryoid bodies using TRIzol (15596-026, Invitrogen), further purified with RNeasy kit (74004, Qiagen) and DNase treatments according to the manufacturer’s instructions. Isolated RNA (1 μg per sample) was used to program cDNA synthesis (SuperScript™ III One-Step RT kit, 12574-018 Invitrogen). Expression of pluripotency markers and determination of tri-lineage differentiation potential was evaluated using the TaqMan hPSC Scorecard™ Panel (A15876, Applied Biosystems) with StepOne Plus™ Real-Time PCR System (4376598, Applied Biosystems). Gene expression data was analysed via the web-based hPSC Scorecard™ analysis software.

### Caridomyocyte differentiation

For generating cardiomyocytes from hiPSC cells cultured on Geltrex or modified Caf1 scaffolds, coated 60 mm culture dishes were seeded at a density of 9 × 10^3^ cells/cm^2^ and cultured as outlined previously. After two or three days, when the colony confluency reached 60–70% cells were treated with 10 μM CHIR-99021 (4423, Tocris Bioscience) and 10 ng/mL recombinant Activin-A (338-AC, Bio-Techne) in RPMI 1640 (61870036, Gibco) with added B-27 supplemented minus insulin (12587010, Gibco). After 24 hours the medium was changed for RPMI-B27 without insulin only and cultured for 48 hours. On day 3 the culture was treated with 5 μM IWP2 (3533, Tocris Bioscience) diluted in RPMI-B27 minus insulin and incubated for a further 48 hours. On day 5 the media was again changed to RPMI-B27 without insulin only and cultured for another 48 hours. At day 7 culture media was changed to RPMI supplemented with B27 containing insulin (17504044, Gibco), no further media supplementations from this point and media replenished every two days thereafter. See **Fig. 5A** for a schematic representation of the protocol.

### Statistical analysis

Data analysis was performed using GraphPad Prism version 9.5.1. for Windows (GraphPad Software LLC, CA, USA). iPSC seeding test was measured on cell counts and analysed using one-way ANOVA, followed by Bonferroni’s multiple comparisons tests. Gene expression was measured using relative quantification and significance between sets determined by one way ANOVA coupled with Šídák’s correction. Statistical significance was considered when *p* <0.05.

## Author Contributions

Conceptualization: ARC, DTP, HW, JHL, RNL. Methodology: ARC, DTP, HW. Formal analysis: ARC. Investigation: ARC, DTP, HW. Resources: HW. Writing—original draft preparation: DTP. Writing—review and editing: ARC, DTP, JHL, RNL, ZMC-L. Visualization: ARC, DTP. Supervision: JHL, RNL, ZMC-L. Project administration: JHL, RNL, ZMC-L. Funding acquisition: JHL, RNL, ZMC-L.

